# A unified framework for geneset network analysis

**DOI:** 10.1101/699926

**Authors:** Viola Fanfani, Giovanni Stracquadanio

## Abstract

Gene and protein interaction experiments provide unique opportunities to study their wiring in a cell. Integrating this information with high-throughput functional genomics data can help identifying networks associated with complex diseases and phenotypes.

Here we propose a unified statistical framework to test network properties of single and multiple genesets. We focused on testing whether a geneset exhibits network properties and if two genesets are strongly interacting with each other.

We then assessed power and false discovery rate of the proposed tests, showing that tests based on a probabilistic model of gene and protein interaction are the most robust.

We implemented our tests in an open-source framework, called Python Geneset Network Analysis (PyGNA), which provides an integrated environment for network studies. While most available tools are designed as web applications, we designed PyGNA to be easily integrated into existing high-performance data analysis pipelines.

Our software is available on GitHub (http://github.com/stracquadaniolab/pygna) and can be easily installed from PyPi or anaconda.

## 1 Introduction

The availability of high-throughput technologies enables the characterization of cells with unprecedented resolution, ranging from the identification of single nucleotide mutations to the quantification of protein abundance [1]. However, these experiments provide information about genes and proteins in isolation, whereas most biological functions and phenotypes are the result of interactions between them. Protein and gene interaction information are becoming rapidly available thanks to high-throughput screens [2], such as the yeast two hybrid system, and downstream annotation and sharing in public databases [3, 4]. Thus, it is becoming obvious to use interaction data to map single gene information to biological pathways.

Integrating interaction information with high-throughput experiments has proven challenging. The vast majority of existing methods are based on the concept of over-representation of a candidate set of genes in expert curated pathways [5, 6]; however, this approach is strongly biased by the richer-get-richer effect, where intensively studied genes are more likely to be associated with a pathway [7], thus limiting the power of new discoveries. Many methods have now been proposed to directly integrate network information for function prediction [8, 9, 10], module detection [11], gene prioritization[12] and structure recognition[13]. However, the vast majority of this software comes either as a web application, visualization plugins or are limited to the characterization of few network properties [14]; while these are simple to use for targeted analyses, they are also difficult to integrate in high-throughput data analyses pipelines.

We believe that, with the availability of big interaction resources and standardized high-throughput analysis pipelines, having a unified and easy to use framework for network characterization of genes and proteins will generate useful information for downstream experimental validation.

Here we build on recent advances in network theory to provide a unified statistical framework to assess whether a set of candidate genes (or geneset) form a pathway, that is genes strongly interacting with each other. We then extended this framework to perform comparisons between two genesets to find similarities with other annotated networks, as a way to infer function and comorbidities. We called our statistical tests geneset network topology test (GNT) and geneset network association test (GNA), respectively. We defined these tests with respect to interactions following a shortest path and random walk model [15].

We then performed extensive analyses to assess power and false discovery rate of our tests; we show that our random walk base statistics are sufficiently powered to detect networks and robust to false discoveries induced by noisy genesets.

We implemented our tests into a Python package, called Python Gene Network Analysis (PyGNA). Our software is designed with modularity in mind and to take advantage of multi-core processing available in most high-performance computing facilities. PyGNA facilitates the integration with workflow systems, such as Snakemake [16], thus lowering the barrier to introduce network analysis in existing pipelines. PyGNA is released as an open-source software under the MIT license; source code is available on GitHub (http://github.com/stracquadaniolab/pygna) and can be installed either through the PIP utility or the anaconda installer.

## 2 Materials and methods

Let *G* = (*V, E*) be a network, or graph, with |*V*| nodes and |*E*| edges. Let *A* be a matrix |*V*| × |*V*|, with *A*_*ij*_ = 1 if there is an edge between node *i* and *j* and 0 otherwise; we denote *A* as the adjacency matrix of the network *G*. We hereby consider only undirected graphs, thus *A*_*ij*_ = *A*_*ji*_. In this context, nodes represent genes or proteins, whereas edges the intervening interactions, e.g. physical, genetic interactions.

Let *S* = *s*_1_,…, *s*_*n*_ be a geneset consisting of *n* genes, we want to quantify the amount of interaction between genes in the geneset (geneset network topology, GNT), and the amount of interactions with genes in another geneset (geneset network association, GNA).

We assume that the strength of interaction between any two genes is a function of their distance on a network *G*, that is closer genes are more likely to interact. Thus, we denote with *i → j* a path in *G* from node *i* to node *j*, whose length, *l*_*ij*_, is the number of edges from *i* to *j*. It is important to note that the concept of path length is well defined only for connected networks, that is ∀_*i, j*_ ∈ *V*, ∃*i* → *j*. We can characterize the strength of interaction between two genes, *i, j*, as the length of the shortest path from *i* to *j*, denoted as *s*_*ij*_. While modeling gene interactions based on shortest path provides a simple and computationally efficient analysis framework, this approach can lead to false positives because biological networks show small-world properties, that is every node is reachable by any other node in the network in few steps [17].

Thus, we introduce a probabilistic model of gene interactions that considers all possible paths between nodes. Let *W* be a stochastic matrix inferred from the adjacency matrix *A*, the probability of reaching node *i* from node *j* after *k* steps is (*W*^*k*^)_*ij*_; this expression describes a random walk (RW) over a network [15]. However, for *k* big enough, the probability of interaction between nodes converges to a value proportional to the degree of the nodes, thus loosing any local structure information. We instead consider a random walk with restart model (RWR), where it is possible to return to the starting node with fixed probability *β*. Importantly, we can estimate the probability of interaction at steady state without losing local structure information as follows:

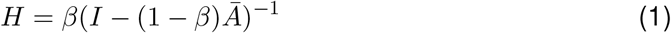

where 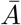 is the normalized adjacency matrix obtained as 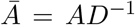, with *D* being the diagonal matrix of node degrees. Under this formulation, the matrix *H* can be interpreted as the heat exchanged between each node of the network, such that closer nodes exchange more heat than distant ones [11]. The RWR model is more robust than the shortest path model, because it effectively adjusts for network structure; it rewards nodes connected with many shortest paths, and penalizes those that are connected only by path going through high degree nodes.

We hereby define both the GNT and GNA tests under a shortest path and RWR model and discuss their interpretation.

### 2.1 Geneset network test statistics

Let *S* = *s*_1_,…, *s*_*n*_ be a geneset of *n* genes, each mapped to a node in *G* = (*V, E*). We are interested in testing whether the strength of interaction between nodes of the geneset is higher than expected by chance for a geneset of the same size.

Under a shortest path model, we define the test statistic *T*_*SP*_ for the geneset *S* as follows:

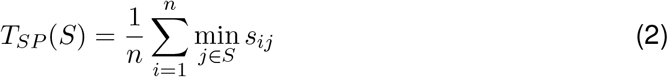

which is the average of the minimum distance between each gene and the rest of those in *S* [18]. Conversely, under a RWR model, we can consider *h*_*ij*_ ∈ *H* as the heat transferred from node *i* to node *j*, which can be used as a measure of interaction strength between the nodes in the geneset *S*, as follows:

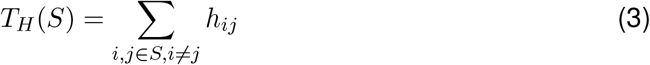

Intuitively, low values of *T*_*SP*_ or high values of *T*_*H*_ are indicative of genes strongly interacting with each other, a phenomenon we call network effect.

Let *S*_1_ and *S*_2_ be two geneset with *n* and *m* genes respectively, we want to estimate the association between *S*_1_ and *S*_2_ as a function of the strength of interaction between their nodes. Under a shortest path model, the association statistics *U*_*SP*_ is defined as follows:

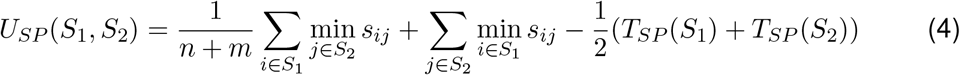

wheareas, under a RWR model, we measure association as a function of the heat, *U*_*H*_, transferred between the two genesets as follows:

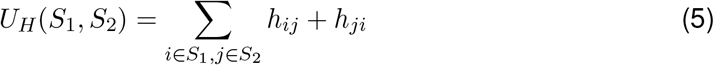

where we consider also the heat withhold by a gene, in case there were overlapping genes between *S*_1_ and *S*_2_.

It is straightforward to prove that shortest path statistics are prone to small-world effects, because if a gene is directly connected with a least one other gene in the geneset, the minimum distance will be 1 regardless of the distance from the other genes in the geneset. Instead, RWR statistics are more robust because they effectively averages the probability of interaction between two nodes with respect to all possible paths. Therefore, we say that RWR statistics have global network awareness, whereas shortest path statistics have local network awareness. It is also important to note that a shortest path model of interaction is also sensitive to missing interactions, which is a well known problem mostly due to experimental limitations. Conversely, a RWR model is able to account for interactions that are likely to occur but difficult to detect [19].

### 2.2 Hypothesis testing

The topological and association statistics are ultimately used for hypothesis testing. To do that, we need a calibrated null distribution to estimate whether the observed statistics are more extreme than what expected by chance. Closed form definition of null distributions is possible only for very simple network models, which are often unrealistic. Therefore, we reverted to a bootstrap procedure to estimate null distributions of the test statistics, conditioned on the geneset size; while this approach can be computationally taxing, in practice, we observed that ≈ 500 bootstrap samples are sufficient to obtain a stable distribution.

Thus, without loss of generality, let *Q* be the null distribution of the test statistic *q* estimated for a geneset of size *n*, and 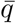 the observed value. It is possible to derive an empirical p-value as follows:

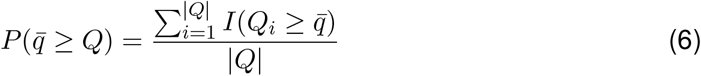

where *I* is the indicator function returning 1 iff the evaluated condition is true. It is straightforward to adapt this formula to the case of testing whether a test statistic is smaller than expected by chance.

For the GNA tests, it is important to note that we are now dealing with two genesets. Hence, a null distribution can be computed either by sampling two random genesets or by sampling only one of the two; we recognize that the latter is more conservative, and is recommended when checking for association with known pathways (see Supplementary Materials).

### 2.3 Implementation

We implemented our statistical tests as part of the Python Gene Network Analysis (PyGNA) framework. PyGNA is implemented as Python package, which can be used as a standalone, command-line application or as a library to develop custom analyses. It is important to note that PyGNA does not provide only GNT and GNA tests, but also integrates standard tests for common network metrics, including internal node degree and module size. PyGNA effectively provides an integrated solution for network analysis (see Supplementary Materials).

Our software can read and write genesets either in Tab Separated Value (TSV) format, Gene Matrix Transposed (GMT), and DESeq2 results in Comma Separated Value (CSV) format [20]. Interaction data can be imported using standard Tab Separated Values (TSV) files, with each row defining an interaction.

Performing tests on large networks using either shortest path or random walk models is computationally taxing. However, since the node pairwise metrics are dependent only on the network structure, they can be computed upfront as part of a pre-processing step. Moreover, since the resulting matrices are usually large and dense, we save them in Hierarchical Data Format (HDF5) format, using the pytables framework [21]. PyGNA performs efficiently both on low-memory machines, using memory mapped input output, and high-performance computing environments, by loading matrices directly in memory.

A bottleneck of the analysis is the bootstrap procedure used to obtain a null distribution for hypothesis testing. However, generating bootstrap samples is a seamlessly parallelizable process, since samples are independent; thus, we implemented a parallel sampler using the multiprocessing Python library, allowing the user to set the number of cores to use. If only one core is requested, the multiprocessing architecture is not set-up, sparing the overhead incurred by setting up a scheduler for running only one thread (see Supplementary Materials).

PyGNA can produce publication ready plots for the different tests, and export network and genesets in standard network formats compatible with graph visualization software, such as Cytoscape [22].

### 2.4 Simulated network and geneset data

We assessed the performance of our GNT tests with respect to different network and geneset. Here, we defined two models encoding different assumptions about network structure and connectivity. First, we generated networks using a stochastic block model (SBM), which allow us to create networks with a controllable number of clusters and probability of interaction. In our context, SBM models are useful positive controls to assess the power of a test in detecting a network effect (see Supplementary Materials).

However, biological networks are poorly described by a simple SBM model, and have properties consistent with experimental and biological confounders [23]. In particular, it has been shown that genes that are intensively studied have more reported interactions (see Supplementary Materials). While this might be explained by a gene controlling multiple functions, a geneset made of a large proportion of high-degree genes does not necessarily represent a network; for example, complex disease genesets have shown limited evidence of co-localization, suggesting that multiple sub-networks could exist within a geneset [24]. Moreover, the presence of high-degree nodes could bias the statistics, since they are central nodes of a network and thus more likely to be connected with other genes. Thus, we reasoned that a robust test should be able to distinguish real modules of strongly interacting genes from the case where high degree genes disrupt multiple, not necessarily interacting, small modules of other genes. To do that, we created networks with nodes connected to each other with probability, *p*_0_, whereas a subset, which we denoted as high degree nodes (HDN), are connected to other nodes with probability *p*_*H*_ > *p*_0_. After the network is generated, we created genesets using a mixture of HDNs and regular nodes to model 3 different scenarios (see Fig. 1): i) partial genesets, consisting of a subset of HDNs; ii) extended genesets, consisting of a subset of HDNs and other randomly selected nodes; iii) branching genesets, consisting of as subset of HDNs, which are expanded with random nodes visited by a depth first search of fixed length and by avoiding loops. The partial and extended genesets are parametrized with respect to percentage of HDNs to include in the geneset whereas, for the extended genesets, we also specify the fraction of random nodes to add as a function of the number of HDNs (see Supplementary Materials). Taken together, we tested our methods on 5790 genesets on a total of 330 networks.

**Figure 1:**
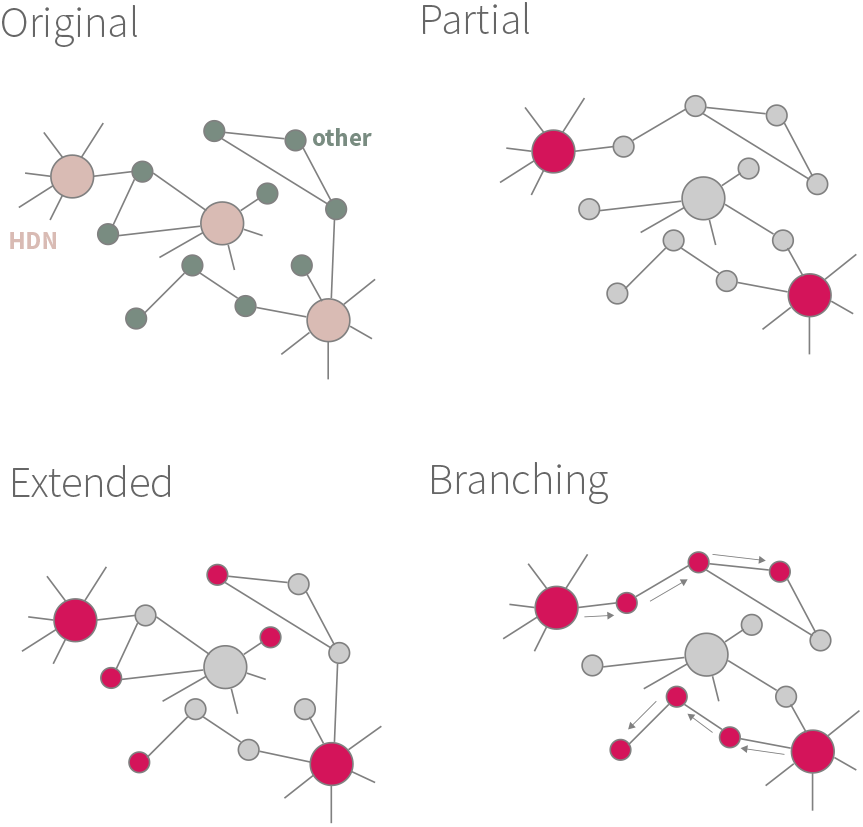
Graphical representation of the generative processes underlying high-degree genesets construction. The original graph is a representation on the input network structure, where only few high degree nodes (HDNs) are present. For each model, red nodes are those assigned to a geneset.

For assessing GNA performances, instead, we created simulated genesets starting from the Gene Ontology (GO) biological process (BP) terms. Network analysis encodes knowledge about gene wiring, hence we assessed whether a GNA test can recognize a geneset made of genes annotated with a GO term and their neighbors. The new genesets were created by randomly sampling genes from a GO term and then extending this set with their neighbors at random; by testing each geneset against its original GO term, we can estimate the power of our test. In our experiments, we parametrized the size of the simulated genesets with respect to the proportion of annotated and neighboring genes.

### 2.5 Real networks and geneset data

We focused on two well characterized interaction datasets: the first is a comprehensive network that aggregates interaction data obtained from 7 different expert curated sources, which we refer to as the interactome [18]; the second is a network defined by physical and genetics interactions reported in BioGRID [3]. Standard metrics were computed for both networks and reported in Tab. 1. It is worth noting, that the interactome had been created using a previous release of BioGRID, and includes also interactions inferred from KEGG pathways. The BioGRID dataset instead is a collection of interactions observed experimentally, thus less prone to study biases.

**Table 1:** Network properties. For each interaction dataset, we report the number of nodes and edges, the density, the median and max node degree (D), and the size of the largest connected component (LCC).

| Network | Nodes | Edges | Density | median(D) | max(D) | LCC |
| --- | --- | --- | --- | --- | --- | --- |
| BioGRID | 17331 | 283991 | 0.00189 | 14 | 2349 | 17310 |
| Interactome | 13460 | 138427 | 0.00152 | 7 | 870 | 13329 |

We used two different geneset datasets in our analyses; the first released as part of the interactome study [18] and the other available from the DisGeNET project [25]. Both the sources have been organised using the MEdical Subject Headings (MESH)^1^. Descriptive statistics for the genesets are reported in Tab. 2. In our experiments, we also used as control genesets the KEGG pathways and the Gene Ontology (GO) terms available from MigSigDB (http://software.broadinstitute.org/gsea/msigdb/).

**Table 2:** Genesets properties. For each dataset, we report the number of geneset, and the minimum, median and maximum number of genes (*n*). The DisGeNET cancer dataset was filtered to keep only genesets with at least 20 genes.

| Dataset | Genesets | min($n$) | median($n$) | max($n$) |
| --- | --- | --- | --- | --- |
| Interactome | 299 | 1 | 27 | 1126 |
| Interactome, GWAS | 299 | 1 | 19 | 370 |
| DisGeNET | 4360 | 2 | 5 | 2037 |
| DisGeNET, GWAS | 622 | 2 | 11 | 595 |
| DisGeNET, cancer groups* | 39 | 22 | 65 | 601 |

<sup>1</sup>https://www.nlm.nih.gov/mesh/meshhome.html

## 3 Results

We tested our geneset network tests implemented in PyGNA on simulated and real datasets; while simulations provide insights into the performances of each test, the analysis of real datasets allowed us to test which model provides the most useful readout.

### 3.1 Simulated networks and genesets

We built a large set of simulations and performed statistical testing using GNT and GNA tests under a shortest path and RWR model. Using SBM simulations, we were able to assess the power, that is the ability of detecting a network effect, if any. The HDN network models, instead, provide a convenient way to estimate the false discovery rate (FDR) of our tests.

Results on SBM networks show that *T*_*H*_ and *T*_*SP*_ are effective in capturing general properties of the geneset. To do that, we analyzed the power and the z-score transform of the statistics (Fig. 2); we hereby consider all SBM genesets with *θ*_*ii*_ > *θ*_0_ to show a network effect. We found that *T*_*H*_ and *T*_*SP*_ return the same genesets as significant, with sufficient power in all cases. We are indeed expecting to see a transition, when the geneset is more connected within itself than with the rest of the network (*θ*_*ii*_ ≤ 0.5). However, it is interesting to note that, as the density of the network increases, the power of *T*_*SP*_ drops even for genesets with high probability of connection (*θ*_*ii*_ = [0.7, 0.8]). This can be better explained by looking at the z-scores; while the *T*_*H*_ statistics has a wider dynamic range, *T*_*SP*_ z-score saturates around ~ 3, with the transition between significant and non significant genesets hardly identifiable. While we recognize that creating the genesets with a SBM model could favour an RWR test statistic, these results show that the shortest path tests are less sensitive and can fail to recognize clear networks effects.

**Figure 2:**
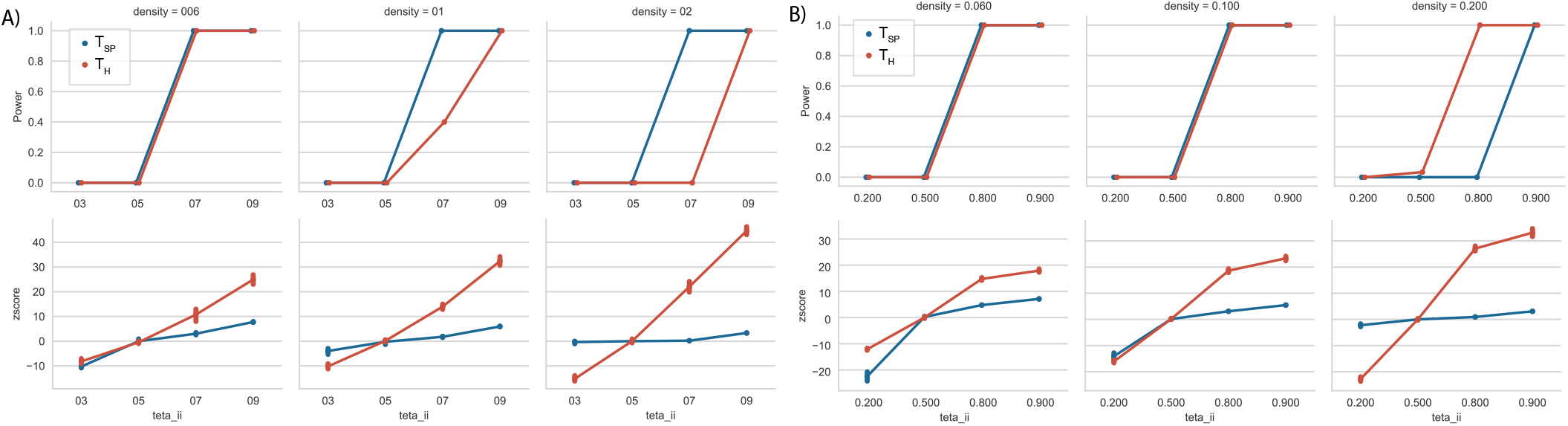
Power analysis of geneset network topological (GNT) tests on networks with a stochastic block model structure. Results are grouped by network density. A) Power and z-score of the 2 block model simulated genesets, at different values of within geneset connection probability, *θ*_*ii*_. B) Power and z-score of the 3 block model simulated genesets, at different values of within geneset connection probability, *θ*_*ii*_.

We then looked at the more realistic networks generated by our HDN model. For the branching datasets, both *T*_*SP*_ and *T*_*H*_ consistently detected a network effect for each geneset. Interestingly, for the branching datasets, when the geneset of interest is made of 2 or more sub-networks, they erroneously detect a network effect. Therefore, we conclude that the presence of multiple sub-networks within a geneset is hardly detectable by topological tests (see Supplementary Materials).

Conversely, the partial and extended datasets highlight different performances for the two tests. In particular, the partial genesets seem to be erroneously called significant by the *T*_*SP*_ test, while *T*_*H*_ is more robust. In particular, 396 genesets are significant for *T*_*SP*_, compared to the 1 from *T*_*H*_. This trend clearly shows that *T*_*SP*_ is prone to false discoveries when HDNs are present in the geneset, although we recognize this to be an unrealistic scenario in practice.

Then, we assessed our test on the extended genesets, which are those more likely to be generated by high-throughput experiments, e.g. RNA sequencing. Here, we consider the genesets to be false networks, since they have been generated by adding an increasing proportion of random nodes to a set of HDN. We then discarded all those with a largest connected component (LCC) containing more than 75% of the genes in the geneset because, despite being random nodes, they show a level of connectedness compatible with a network effect. Then, we estimated a false discovery rate (FDR) for both tests, as the fraction of genesets reported as significant. It is striking that, for a geneset with less than 10 HDNs, the *T*_*H*_ has *FDR <* 10%, whereas *T*_*SP*_ has consistently *FDR >* 10% (Fig. 3). For genesets made of 10 or more HDNs both tests are not reliable; however, in practice, genesets with the same proportion of HDNs are unlikely.

**Figure 3:**
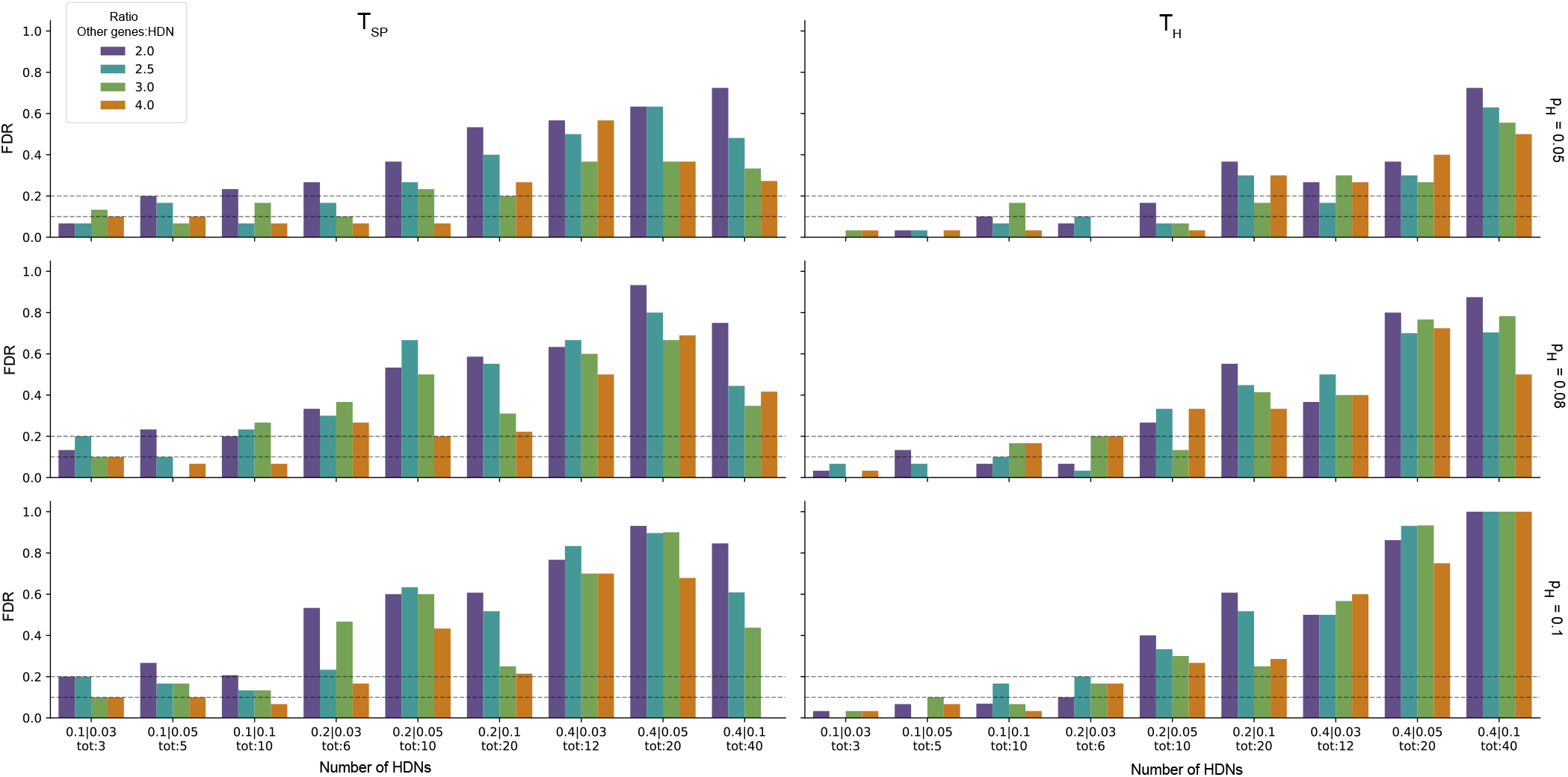
Results for the extended high degree nodes (HDNs) simulations. We report the False Discovery Rate (FDR) of *T*_*SP*_ and *T*_*H*_ for each parameter setting.

For the GNA tests, our simulations mimic the common scenario where a geneset is tested for association with known pathways and GO terms. We then compared our results with those of an over-representation analysis (ORA) via a Fisher’s exact test. It is clear that the *U*_*H*_ test has more power than *U*_*SP*_ and ORA (Fig. 4). As expected, as soon as the number of genes shared between the geneset and the GO drops, the ORA analysis fails to recognize any association between the two geneset; this result provides further evidence that ORA methods are strongly affected by annotation accuracy and thus have limited detection power.

**Figure 4:**
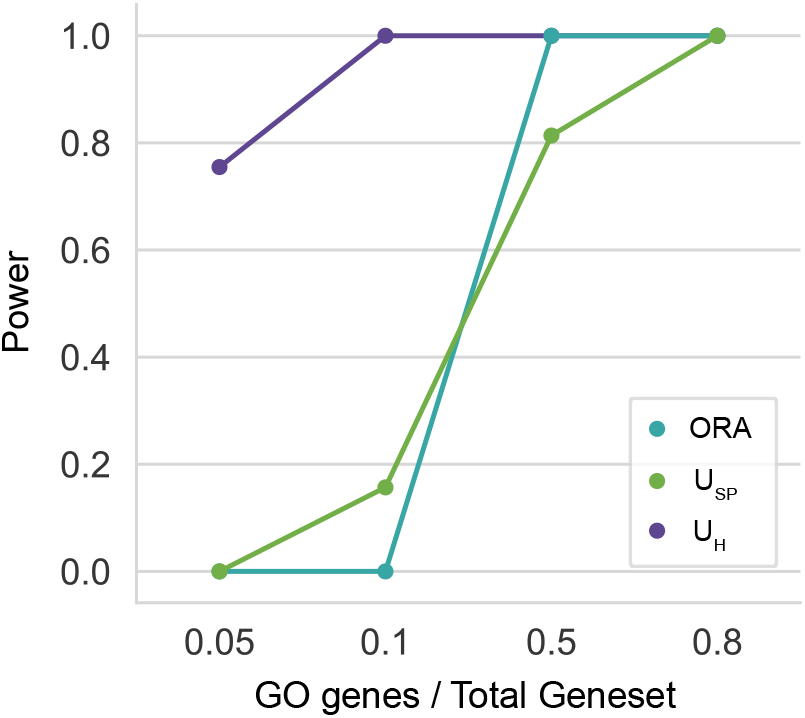
Power results for the geneset network association tests, *U*_*SP*_ and *U*_*H*_, compared to Fisher’s over-representation test (ORA).

### 3.2 Real network and geneset datasets

We then used the GNT tests to analyse real data from two interaction datasets, the interactome and BioGRID, and two geneset databases, the interactome derived and Dis-GeNET. We tested both genesets on the two networks to avoid annotation specific biases.

For most genesets, we have no ground truth on whether they exhibit network properties, thus we decided to use the KEGG pathways as positive controls. We reasoned that since KEGG pathways have been integrated into the interactome, they should show a network effect. We set the threshold for significance at *p* ≤ 0.01, and found all pathways to be significant with the *T*_*H*_ test and only 1 not significant with the *T*_*SP*_ test. We then concluded that both tests are sufficiently powered to detect well organized networks.

We then tested genesets from gene-disease associations databases; despite, we expected the majority to show a network effect, we wanted to test how the two statistics compare and how network structure might influence the results. The *T*_*SP*_ test consistently reports more statistically significant genesets (Fig. 5), finding 95% of the interactome genesets to be significant on the interactome. This result has been suggested to prove that, even in incomplete networks, network topology analysis is able to recognize connected sub-graphs [18]. However, we noticed that the proportion of significant genesets found with the *T*_*H*_ test is remarkably smaller, albeit these genesets have also been discovered with the *T*_*SP*_ test.

**Figure 5:**
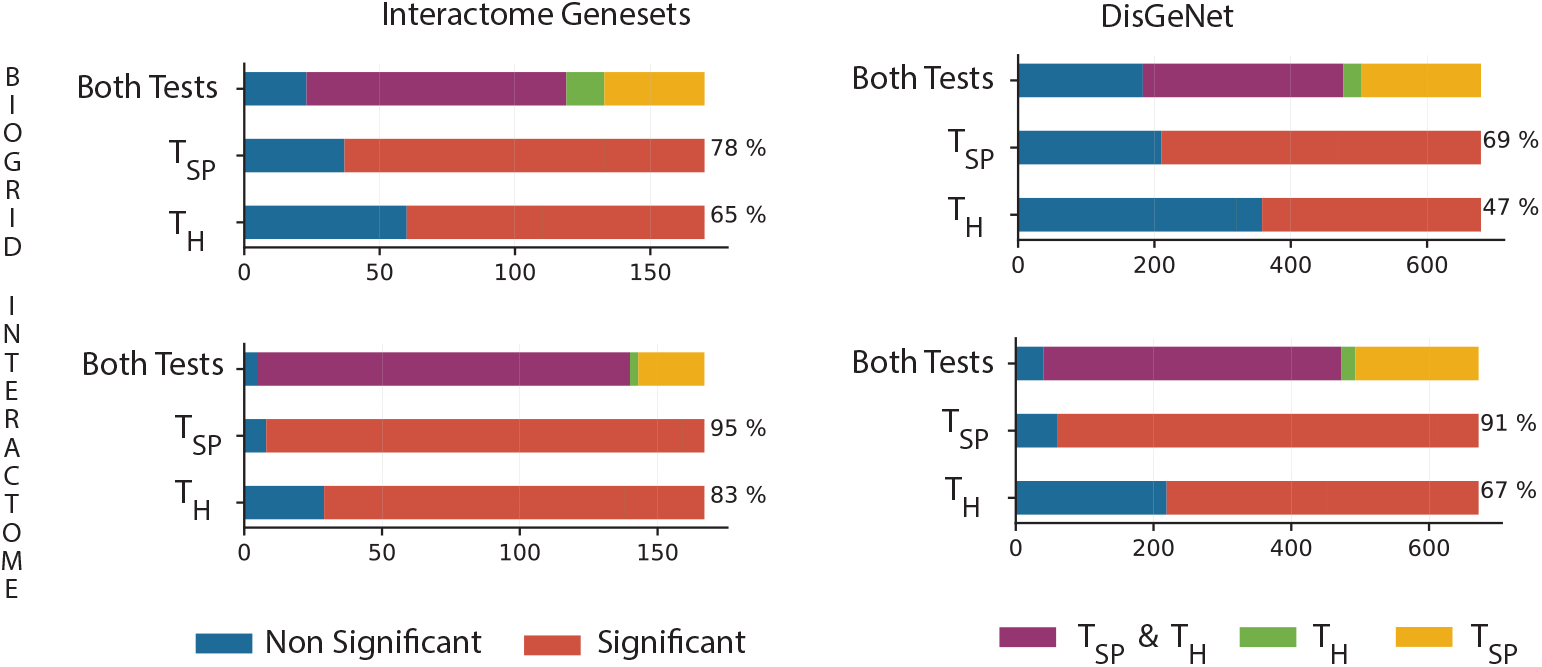
Significant genesets from the gene network topology (GNT) analysis for the interactome and Dis-GeNET genesets across the BioGRID and interactome networks.

Interestingly, we obtained substantially different results when using BioGRID; specifically, we observed a drop of approximately 20% in significant genesets, albeit we saw consensus between *T*_*SP*_ and *T*_*H*_ in the two networks. This result suggests that, for incomplete networks, we can expect a high number of false positives.

Both the interactome and DisGeNET datasets report gene-disease association derived from Genome-Wide Association Studies (GWAS); in general, these genesets are obtained by assigning genome-wide significant single nucleotide polymorphisms (SNPs) to the nearest gene. Since these genesets are usually more representative of the genetic architecture than the molecular mechanisms underpinning a disease, we expected a weaker network effect. Consistent with this hypothesis, when we analyzed the GWAS genesets from both databases, we observed a significant drop in the percentage of significant genesets.

We then used our GNA tests to identify biological processes associated with cancer. It is well-known that cancers acquires specific biological capabilities during tumour formation [26]; thus, we reasoned that cancer genesets should show significant association with many biological processes networks. To do that, we took the 38 cancer genesets, as defined in DisGeNET, and tested for association with 1500 GO biological process (BP) terms, which are those having a number of annotated genes between 50 and 500. Here, we used only our *U*_*H*_ test, since our simulations have shown this test to be consistently the most robust.

Consistent with our hypothesis, we observed well-known biological processes associated with cancer for all genesets; in particular; more than 80% of the genesets showed significant association with cell proliferation, aging, apoptosis and response to oxidative stress processes, which are usually deregulated in tumors (see Supplementary Materials).

We then tested whether biological processes found by GNA could be useful in classifying cancer subtypes. To do that, we performed hierarchical clustering of cancer genesets using the Jaccard distance as metric (Fig. 7); here, for each cancer, we considered a binary vector of GO terms, with 1 if the term is significant (*p* ≤ 0.001), and 0 otherwise. We found that cancers in the same anatomical district (e.g. head and neck, colonic and colorectal) share many genes controlling the same biological processes; as a further evidence, we observed strong clustering between neoplasms often arising in epithelial cells. Thus, we conclude that geneset network association analysis could provide useful information to characterize molecular mechanisms underpinning complex disease phenotypes.

**Figure 6:**
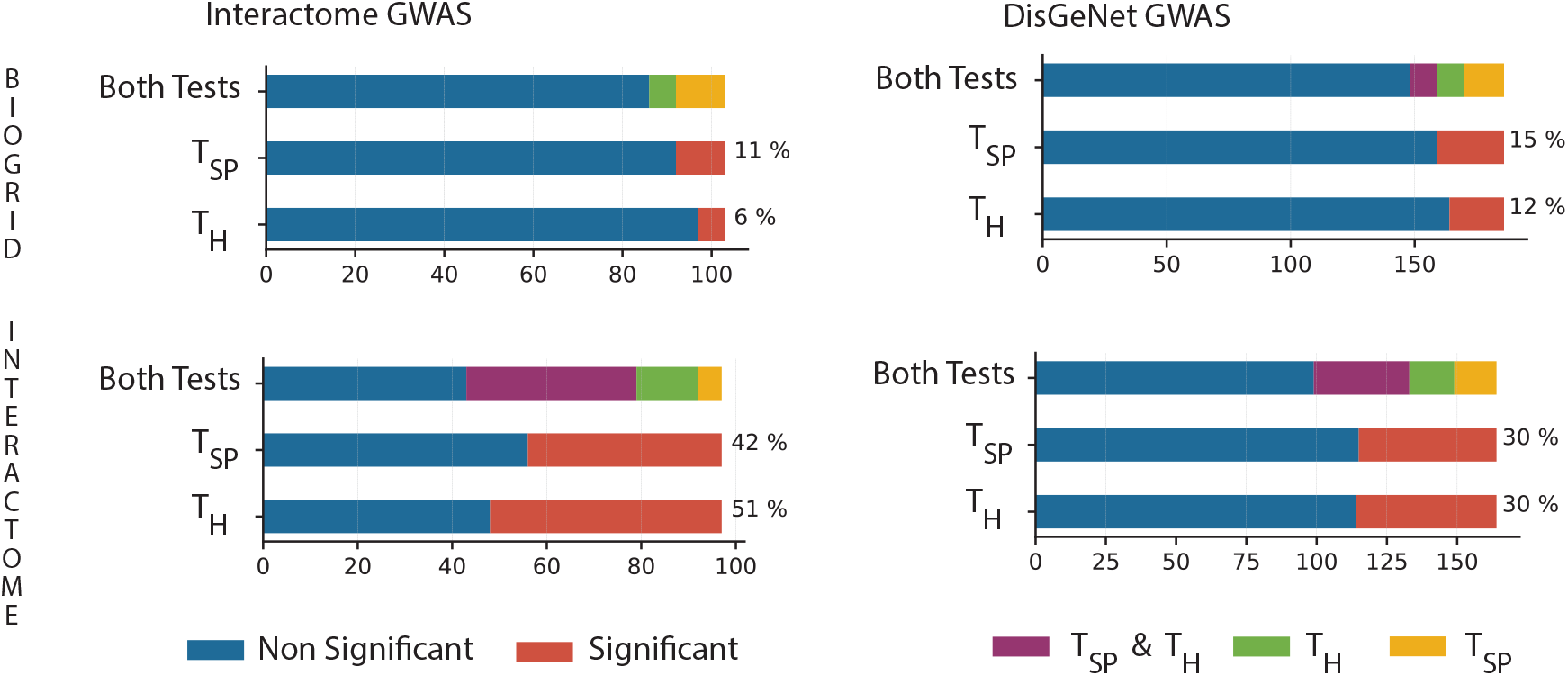
Significant genesets from the gene network topology (GNT) analysis for the interactome and DisGeNET GWAS genesets across the BioGRID and interactome networks.

**Figure 7:**
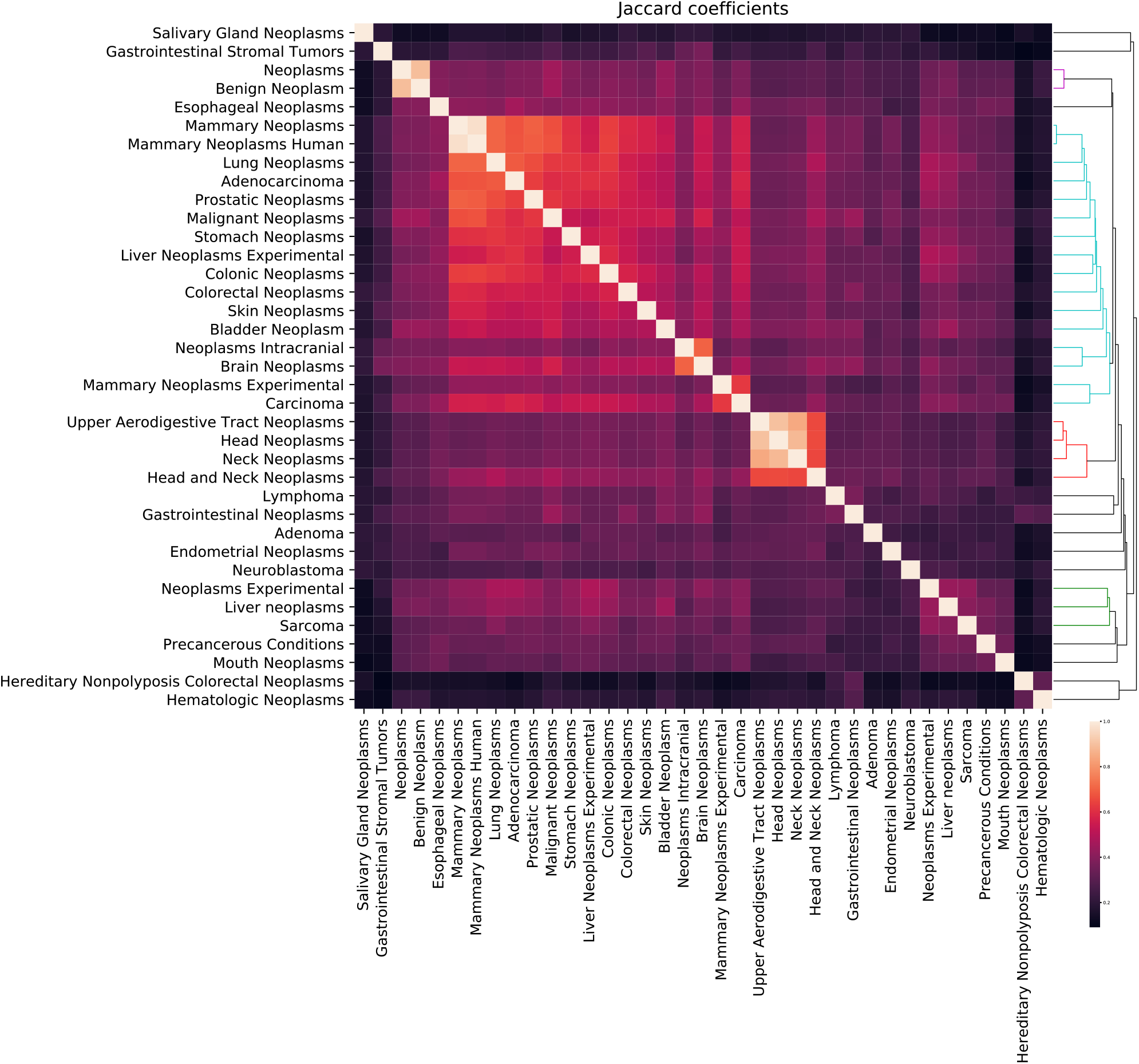
Gene Ontology (GO) hierarchical clustering of 38 different cancer types. The dendogram is obtained by clustering cancers by shared significant GO terms; the heatmap shows the Jaccard coefficient between pairs of cancer groups.

## 4 Discussion

The availability of gene and protein interaction data provide unique opportunities to understand the cellular wiring underpinning most common complex phenotypes. However, integrating network and gene-level information has been challenging.

Here we proposed a unified theoretical and computational framework to study genesets at the network level; we presented statistical tests to study network topology and similarity under a shortest path and random walk model of interaction. We estimated power and false discovery rates under different network and geneset structures, showing that random walk tests are the most robust and should be used for the analysis of noisy data, such as those generated by high-throughput sequencing experiments.

We also recognize the limits of our work. Network topology analysis is sensitive to the network structure; thus it is becoming clear that using up-to-date interaction datasets, which usually define dense networks, is required to obtain robust results. Moreover, given the complexity of gene-to-phenotype associations, topological analysis provides only preliminary evidence of network effects; thus, we anticipate that new statistical tests will be required to capture the inherent complexity of biological pathways.

Our contribution is not only theoretical, but also practical; we implemented our statistical tests as part of a modular Python package, called Python Geneset Network Analysis (PyGNA). Different from existing applications, we designed PyGNA to be easily integrated into workflow systems and rapidly provide a comprehensive network characterization of input genesets. Our software takes advantage of multi-core architectures and can work in both desktop and high-performance computing environments, thus lowering the computational requirements to perform network analysis.

PyGNA is not only a stand-alone application, but also a Python library that can be easily integrated into other software; thus, we envision our framework as an open-source platform to develop network statistical tests.

## 5 Contribution

V.F. and G.S. conceived the study. V.F. wrote the PyGNA and performed experiments under G.S. supervision. V.F. and G.S. wrote the manuscript.

## Footnotes

1 https://www.nlm.nih.gov/mesh/meshhome.html

