## Supplementary Materials for "A unified framework for geneset network analysis"

Viola Fanfani [*], Giovanni Stracquadanio, [†]

July 11, 2019

**2 Geneset network association bootstrap procedures** **2**

**3 Parallel sampling procedure** **2**

**4 Stochastic Block Model Simulations** **4**

**5 Real networks properties** **8**

**6 Results for the high degree nodes branching network model** **10**

**7 Gene ontology network association results for $38$ cancers** **11**

[*] School of Biological Sciences, The University of Edinburgh, Edinburgh EH9 3BF, United Kingdom
[†] Corresponding author. School of Biological Sciences, The University of Edinburgh, Edinburgh EH9 3BF, United Kingdom

### 1 Additional geneset network topology tests

PyGNA integrates many other common tests for the analysis of network in addition to our shortest path and random walk with restart models.

The statistical testing framework is the same described for $T_{SP}$ and $T_H$, where an empirical pvalue is obtained through bootstrap resampling. Here, we provide an overview of other available statistics for a single set of $n$ genes $S = s_1, s_2, \cdots, s_n$.

#### 1.1 Module statistics

Let LCC($S$) the largest connected component induced by geneset $S$. We define the module statistic $T_M$ as follows:

$$T_M = |LCC(S)| \tag{1}$$

#### 1.2 Total degree statistics

Let $D_i$ be the degree of the gene $i$ in geneset $S$, that is the number of edges connecting $s_i$ to other nodes. The total degree statistic is defined as follows:

$$T_{TD} = \frac{1}{n} \sum_{i \in S} D_i \tag{2}$$

#### 1.3 Internal degree statistics

Let $\bar{D}_i$ be the internal degree of gene $i$ in geneset $S$, that is the number of edges connecting $s_i$ with another gene $s_j \in S$. The internal degree statistic is defined as follows:

$$T_{ID} = \frac{1}{n} \sum_{i \in S} \frac{\bar{D}_i}{D_i} \tag{3}$$

### 2 Geneset network association bootstrap procedures

We hereby explain how the null distributions are generated for the geneset network association (GNA) tests, which give an estimate of the strength of interaction between two genesets, $S_1, S_2$. When we generate a null distribution by sampling two random genesets of size equal to $S_1$ and $S_2$, we are performing a test under the null hypothesis of no difference between the strength of association observed for $S_1$ and $S_2$ and any two random genesets of equal size. Conversely, when one of the geneset is a Gene Ontology (GO) or pathway term $R$, it is advisable to be more conservative; here, we resample just the input geneset and keep the term $R$ fixed, such that we perform a test under the null hypothesis that there is no difference in strength of interaction between the input geneset and any other random geneset of the same size with respect to the term $R$.

### 3 Parallel sampling procedure

We implemented a parallel sampler to speed up the generation of null distributions for our tests. We tested how our sampler scales as a function of the number of cores allocated using the interactome, as reference network, and generating genesets by taking random nodes from it. We then performed GNT analyses using both the module and random walk statistics, $T_M$, $T_H$, to test the performances when the statistic is estimated from a large matrix and when is evaluated only from the network structure. We performed our tests on genesets of size $[50, 100, 500]$ and by increasing number of cores $[1, 3, 6, 8]$ and permutations $[500, 1000, 10000]$ (Fig. 1).

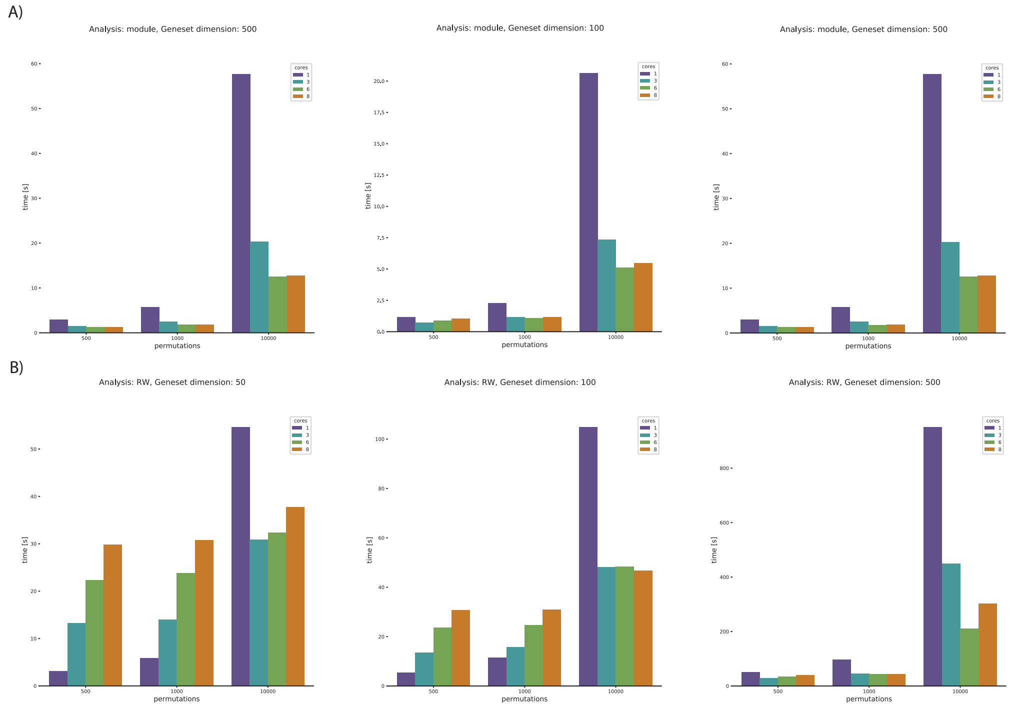

Figure 1: Running time of geneset network topology tests using parallel sampling.

As expected, parallel sampling considerably speeds up the generation of null distributions. However, for small genesets and few permutations, reading the HDF5 file is more demanding than the sampling procedure. It is also worth noting that we ran a test assigning $8$ cores on a machine on a 6-core CPU and, as expected, we observed no improvement.

### 4 Stochastic Block Model Simulations

#### 4.1 Block model simulations

We simulated two and three blocks networks with the genesets being the nodes in each block.

In both cases, we used one of the clusters as background network, containing the vast majority of nodes and having a lower probability of interaction. With these settings, we are able to simulate a large block that is not assortative and one or two clusters that are more connected than the rest of the network by construction.

A stochastic block model with $n$ blocks is described by a membership matrix $Z : n \times n$:

$$Z = \begin{bmatrix} \theta_{11} & \theta_{12} & \dots & \theta_{1n} \\ \theta_{21} & \theta_{22} & \dots & \theta_{2n} \\ \vdots & \vdots & \ddots & \vdots \\ \theta_{n1} & \theta_{n2} & \dots & \theta_{dn} \end{bmatrix} \tag{4}$$

where $\theta_{kk}$ is the probability of having an edge between nodes within the block $k$ and $\theta_{kj}$ is the probability of having an edge between nodes in block $k$ and block $j$. In our simulations, we parametrized the number of nodes of the network $G$, the number of nodes in each block $k$ as a fraction of the total $N_k = p_k N$, the block model matrix $Z$ and the density $d$ of the network, that is a multiplicative term applied to the $Z$ matrix.

For a 2 block network, we used two parameters $p_1$, the percentage of the network nodes $N$ falling belonging to the cluster of interest $C_1$, and $\theta_1$, which is the probability of interaction within the set. With the other cluster $C_0$ having $N(1 - p_1)$ nodes and $\theta_0 = 1 - \theta_1$.

$$Z = \begin{bmatrix} \theta_1 & \theta_0 \\ \theta_0 & \theta_0 \end{bmatrix} \tag{5}$$

In Fig. 2, we are showing how adjacency matrix (restricted only two 200 nodes for better visualization) changes according to the parameters.

For a network with 3 blocks, the parametrization is slightly more complex, but allows to test a broader range of hypotheses. Given the blocks, $C_0$, $C_1$, $C_2$, we parametrise their dimension as $p1 = p2$ with the bigger cluster having $p_0 = N(1 - 2p_1)$ nodes. The membership matrix $Z$ is then parametrized as follows:

$$Z = \begin{bmatrix} \theta_{ii} & \theta_{ij} & \theta_0 \\ \theta_{ij} & \theta_{ii} & \theta_0 \\ \theta_0 & \theta_0 & \theta_0 \end{bmatrix} \tag{6}$$

We assume that the vast majority of nodes are assigned to cluster $C_0$, having fixed probability of interaction $\theta_0$ . To simplify the parametrization, we kept $\theta_{ii} = \theta_{11} = \theta_{22}$ and $\theta_0 = 1 - \theta_{ii}$. The connection between $C_1$ and $C_2$ is $\theta_{ij} = \max(perc_{ij}\theta_{ii}, \theta_0)$, where $perc_{ij}$ modulates the probability of interaction between the two clusters. When $perc_{ij} = 1$ we have a single cluster ($\theta_{11} = \theta_{12} - \theta_{21}, \theta_{22}$). The limit of $\theta$ is given by $\theta_0$, meaning the two clusters are connected with each other as the rest of the network. Overall the parameters are $p_1$, $\theta_ij$ and $perc_{ij}$. In Fig. 3, we visualize the adjacency matrix for different $perc_{ij}$ values.

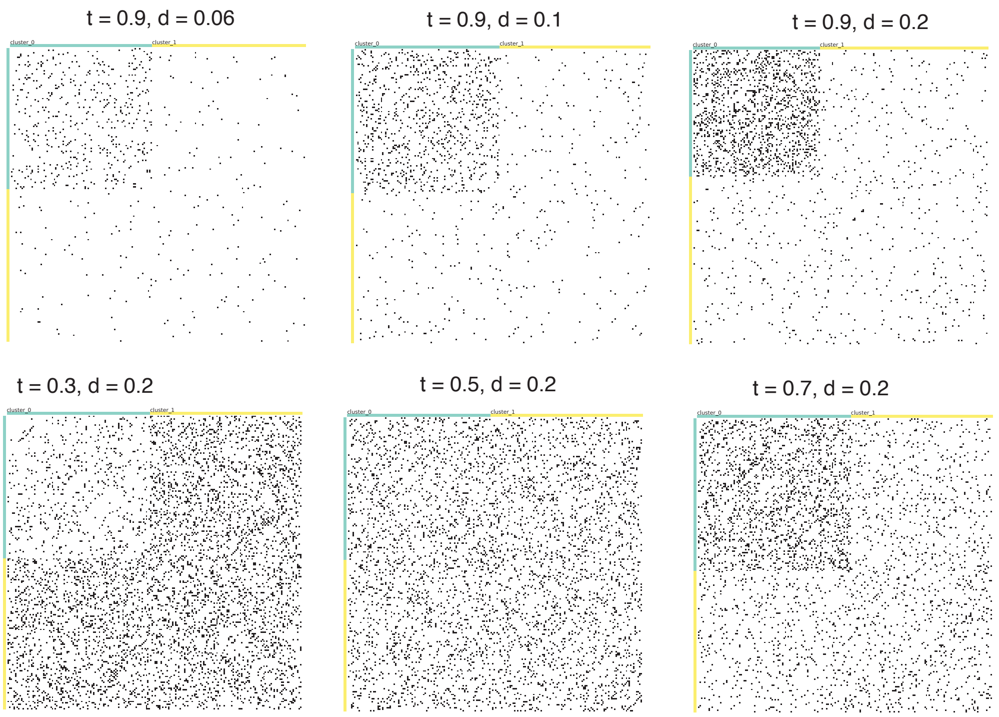

Figure 2: Adjacency matrix from block model simulations with 2 blocks. Here $t = \theta_1$ and $d$ is the density of the network.

###### 4.1.1 Block model parametrisation

The two block simulations have been generated for $N = 1000$ nodes, $d = [0.06, 0.1, 0.2]$ and $p_0 = 0.1, p_1 = 1 - p_0$. The matrix is built by setting $\theta_0 = 1 - \theta_1$ and $\theta_1 = [0.9, 0.7, 0.5, 0.3]$. In Fig. 2, we report the adjacency matrix for different parameters of the block model and network density (only $200$ nodes are reported since the rest of the network is modelled as the large cluster). For each condition we generated $5$ replicates.

As for the three-blocks model, these simulations have been generated for $N = 1000$ nodes and $d = [0.06, 0.1, 0.2]$ $p_0 = p_1$ and $p_2 = 1 - p_0 - p_1$. In Fig. 3, we depicted adjacency matrices for different $perc_{ij}$. For each condition, we generated $5$ replicates. We have shown the results of 3 blocks SBM with parameters: $\theta_{ii} = [0.9, 0.8, 0.5, 0.2]$ and $perc_{ij} = [1, 0.9, 0.5]$.

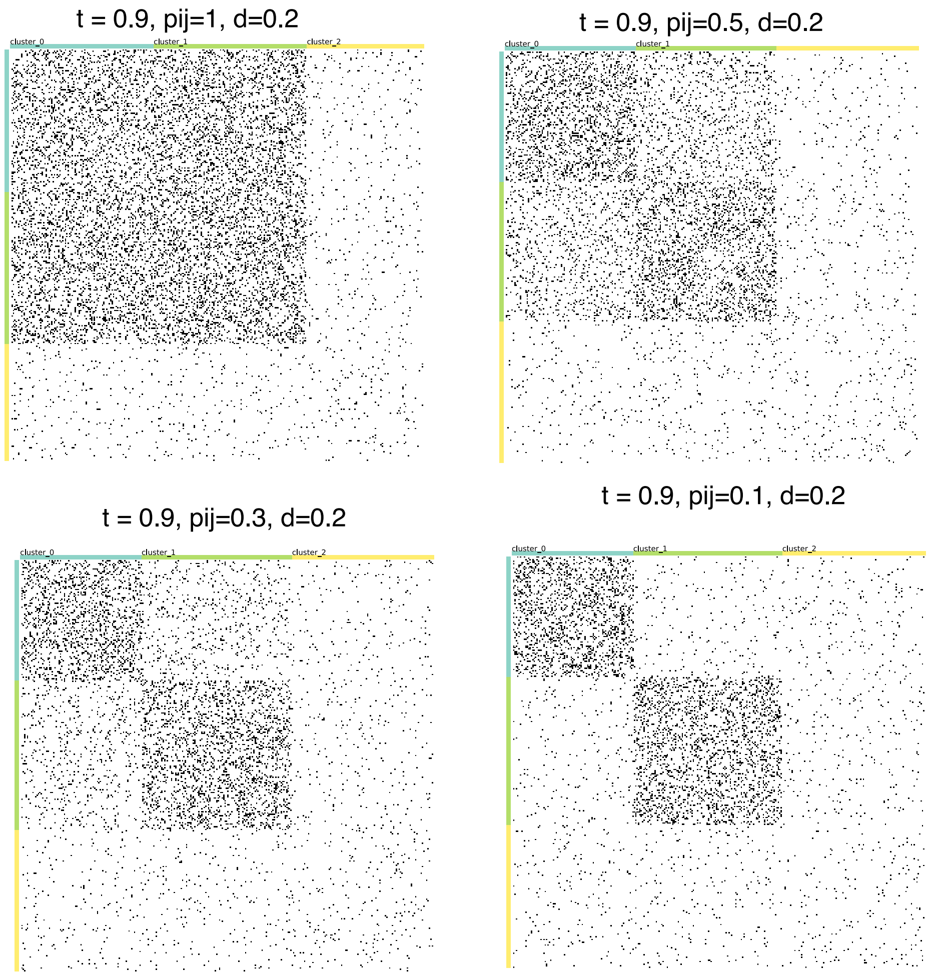

Figure 3: Adjacency matrix for BM simulations with 3 blocks. Here $t = \theta_{11}$ and $d$ is the density of the network.

#### 4.2 High degree nodes networks parametrization

For the high degree nodes (HDN) network simulations, there are two layers of parametrization. First, we defined the properties of the network, including the number of nodes $N$, the probability of connection $p_0, p_H$ for the regular and HDN nodes, respectively. Then, we varied the proportion $perc$ of the total nodes that are HDN. We presented results for $N = 1000$, $p_0 = 0.006$, $p_H = [0.1, 0.08, 0.05]$, $perc = [0.03, 0.05, 0.1]$. For each setting, we generated $10$ networks.

We then selected the geneset to be tested. For the partial genesets, we varied the proportion of HDN to fall into the new geneset, $pHDN = [0.1, 0.3, 0.5]$, with the total number of genes in the geneset being $N \times perc \times pHDN$. For the extended genesets, we specified both $pHDN$ and the ratio between the random nodes added and the HDN in the geneset. Thus, for a new geneset of length $M$ the ratio is defined as $ratio = (M - N \times perc \times pHDN)/N \times perc \times pHDN$ where $(M - N \times perc \times pHDN)$ is the number of new nodes. We showed the results for $pHDN = [0.1, 0.2, 0.4]$ and $ratio = [2, 2.5, 3, 4]$. Finally, for the branching genesets, we used $3, 5$ and $10$ HDN nodes as starting points for generating depth first paths of length $5, 10$ and $15$.

#### 5 Real networks properties

We further characterized our real networks by looking at the node degree distribution. We observed that most nodes have low degree (Fig. 4), albeit the degree distribution has a remarkably long tail.

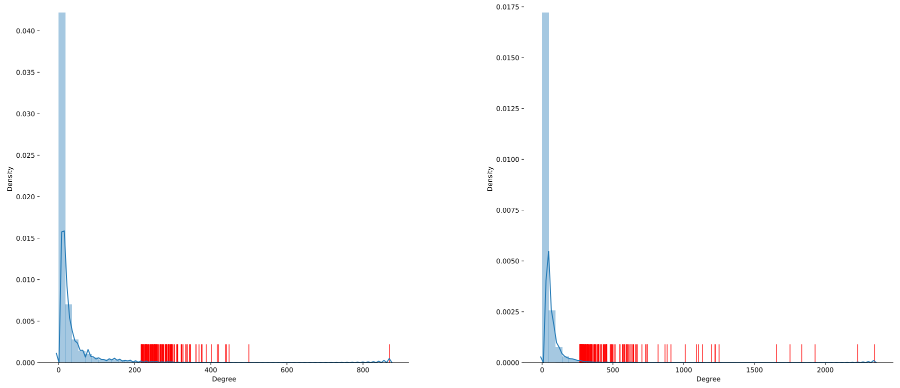

Figure 4: Degree distribution of the interactome and BioGrid networks. We reported in red the genes over the $99$th percentile.

For BioGRID, we were able to study also the number of publications for each reported gene. This way we are able to see if there is some relationship between the degree of the node and the number of studies. As expected, there is correlation between degree and number of studies per gene (Fig. 5), with the vast majority being genes implicated in cancer.

This is a speculative analysis, yet it can give insights into the process of network generation. The fact that some genes are well studied is surely justified by their biological importance. At the same time, we had to account for potential biases introduced by this network property. To do that, we assessed the false discovery rate of our tests using the HDN model.

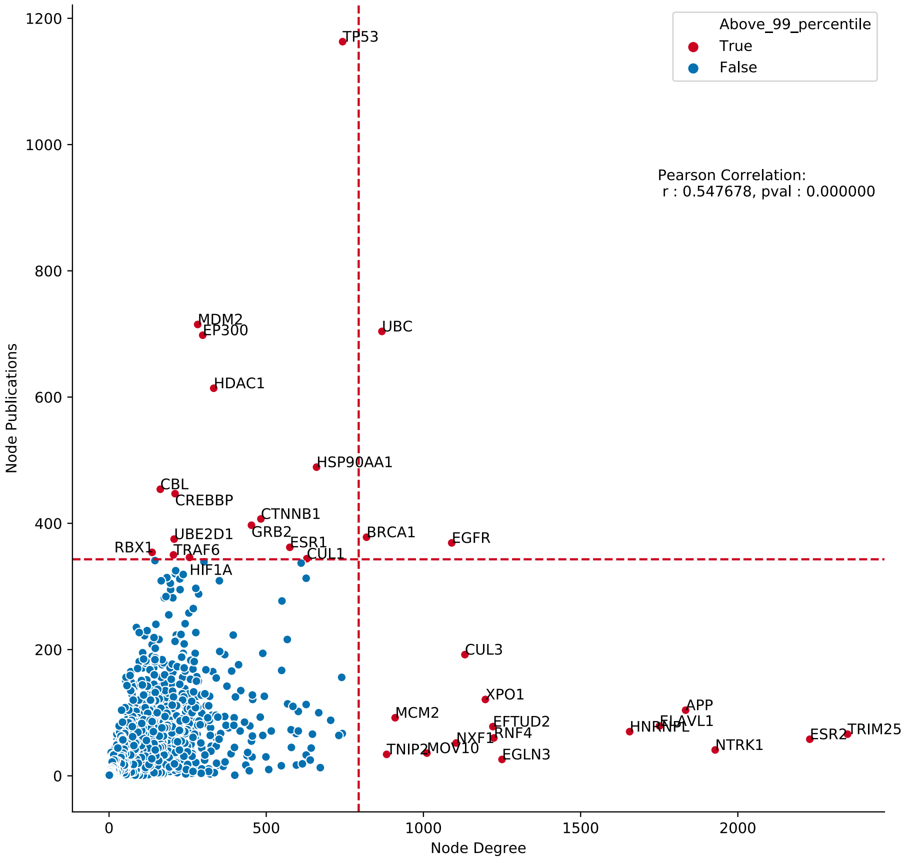

Figure 5: Relationship between the degree and the number of publications for each node in BioGrid.

### 6 Results for the high degree nodes branching network model

The branching genesets have been parametrized using $[3, 5, 10]$ seed HDN nodes to which $[5, 10, 15]$ are attached. From the analysis of the branching genesets, all are reported as significant for both $T_{SP}$ and $T_H$ (Fig. 7).

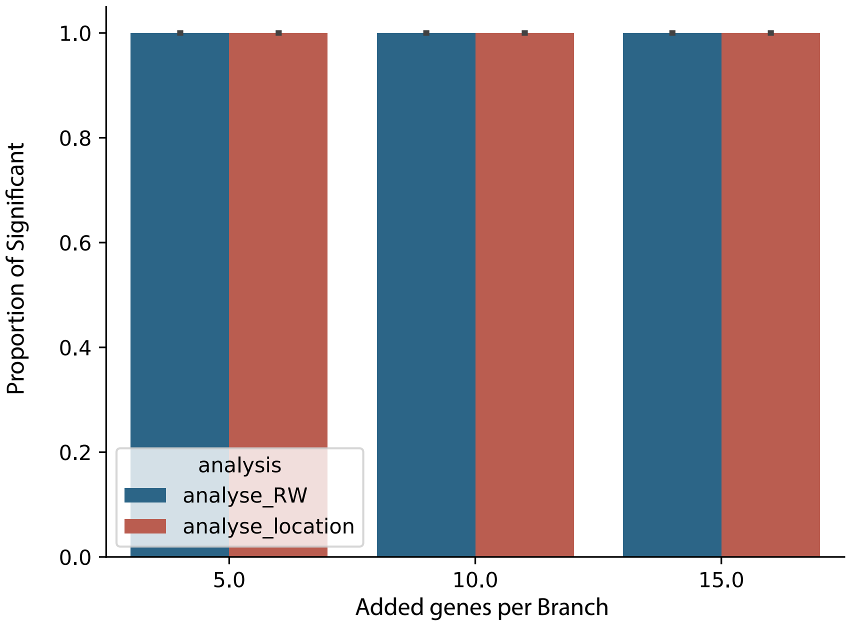

Figure 6: Proportion of significant branching genesets at varying number of new nodes.

### 7 Gene ontology network association results for $38$ cancers

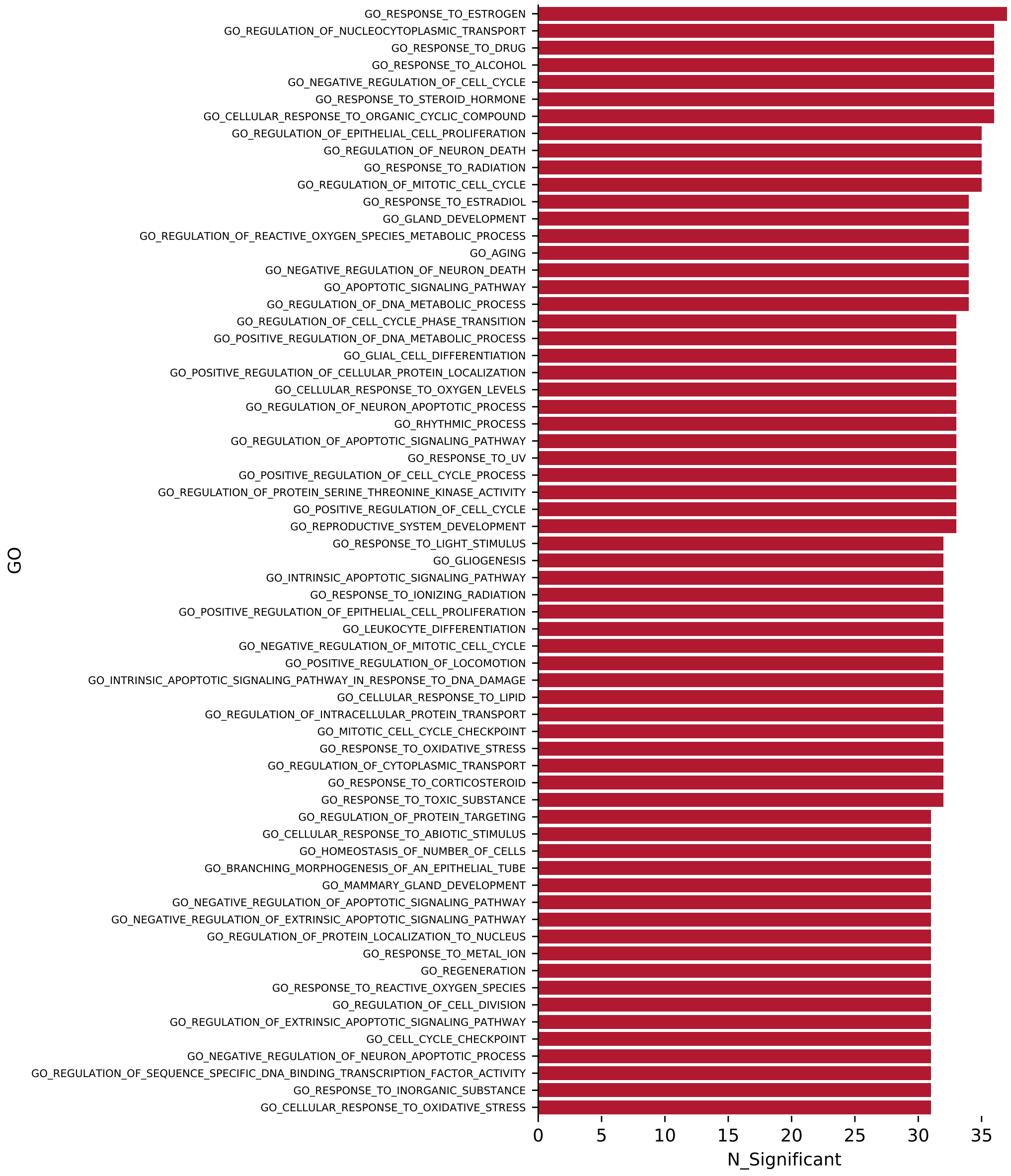

Figure 7: Gene ontology terms associated with multiple cancers at $p \leq 0.001$.
